# EmpiReS: Differential Analysis of Gene Expression and Alternative Splicing

**DOI:** 10.1101/2020.08.23.234237

**Authors:** Gergely Csaba, Evi Berchtold, Armin Hadziahmetovic, Markus Gruber, Constantin Ammar, Ralf Zimmer

## Abstract

While absolute quantification is challenging in high-throughput measurements, changes of features between conditions can often be determined with high precision. Therefore, analysis of fold changes is the standard method, but often, a doubly differential analysis of changes of changes is required. Differential alternative splicing is an example of a doubly differential analysis, i.e. fold changes between conditions for different isoforms of a gene. EmpiRe is a quantitative approach for various kinds of omics data based on fold changes for appropriate features of biological objects. Empirical error distributions for these fold changes are estimated from Replicate measurements and used to quantify feature fold changes and their directions. We assess the performance of EmpiRe to detect differentially expressed genes applied to RNA-Seq using simulated data. It achieved higher precision than established tools at nearly the same recall level. Furthermore, we assess the detection of alternatively Spliced genes via changes of isoform fold changes (EmpiReS) on distribution-free simulations and experimentally validated splicing events. EmpiReS achieves the best precision-recall values for simulations based on different biological datasets. We propose EmpiRe(S) as a general, quantitative and fast approach with high reliability and an excellent trade-off between sensitivity and precision in (doubly) differential analyses.

## INTRODUCTION

Alternative splicing is an important regulatory mechanism that contributes to the complexity of the transcriptomic landscape in higher eukaryotes. More than 85% of the human protein coding genes (GRCh38.95) can generate two or more transcripts. The 3D-structure of a protein can be heavily altered by alternative splicing (1) which may have major effects on the function and phenotype (2). Alternative splicing can also act as an additional regulatory element by producing defunctional transcripts which are degraded prior to translation. Different transcripts of the same gene are often expressed at the same time, possibly at different abundance, yielding a defined mixture of isoforms. Changes to this composition are further referred to as differential alternative splicing (DAS). Various diseases can be linked to DAS events (3). Most hallmarks of cancer like tumor progression and immune escape (4, 5) can be attributed to changes in the transcript composition (6). DAS plays a prominent role in various types of muscular dystrophy (MD) (7), splicing intervention by targeted exon removal is a therapeutic option in Duchenne MD (8). This renders DAS a potential target for novel drug therapies. Alternative isoform regulations (9) also play a crucial role for evolutionary differences between species as analyzed for a range of vertebrate tissues and organs (10, 11).

The current method of choice to investigate DAS as well as changes to total gene expression is mRNA sequencing (RNA-seq). Improvements of sequencing instruments and new techniques have facilitated sequencing depths and qualities that allow to identify differences in the expression on the transcript level on a genome-wide scale. Fortunately, it has also become common to sequence many replicates of the same condition or phenotype, which allows to account for biological variance, bias and measurement noise, a step which is necessary to draw biologically meaningful conclusions. Absolute quantification of transcripts from raw RNA-seq data is still challenging. Many reads map to multiple transcripts. Different transcripts are subject to different biases like length or GC content. Tools like Kallisto (12) or StringTie (13, 14) assign reads to their transcript of origin and try to resolve such ambiguities using expectation-maximization methods. Other tools like Sleuth (15) and Ballgown (16) try to detect transcripts which are expressed differentially between groups of samples. They partially correct for this mapping uncertainty. However, they only allow differential tests, i.e. whether the expression of a transcript changes between two groups of samples. They do not allow to check whether two transcripts change to a different extent or in different directions (doubly differential test), which is necessary to detect DAS. Other tools which explicitly test DAS use these estimated counts and do not account for their uncertainty (DRIMSeq (17) or SUPPA2 (18)). Count based methods for DAS such as DEXSeq (19) or rMATS (20) perform their test on smaller sub-units like exon bins or certain splice junctions which can be directly observed in the data. Typically, reads that overlap with more than one unit are counted multiple times. Furthermore, it is not straightforward to draw conclusions on the transcript level using exon bins.

With MS-EmpiRe (21) we introduced a new approach that does not assume a given distribution for the detection of differentially expressed proteins in mass spectrometry data. Differences between expression on both gene and transcript level are usually estimated in terms of fold changes. For properly defined fold changes, e.g. fold changes for the same objects between different samples, biases which complicate absolute quantification usually cancel out. In that work, we proposed to use empirical and intensity-dependent fold change error distributions. MS-EmpiRe outperformed state-of-the-art tools, yielding highly sensitive results while still properly controlling the number of false discoveries. Here, we extend this approach to EmpiRe to take condition-dependent error distributions into account and show that it is applicable to count-based RNA-seq data and can compete with other tools. Furthermore, we propose EmpiReS which extends this model such that it can be applied to detect “changes of changes” as is necessary for DAS.

To benchmark EmpiReS we use extensive simulated data based on real measurements. In contrast to other benchmark approaches, we do not assume that the counts follow a given statistical distribution, but directly use the counts observed in an experiment. The simulated data represents many features of real data and, thus, will provide a standardized test set to assess a number of methods compared here, but it is also made available to be used for future studies. A similar benchmark is also provided to assess the performance of EmpiRe for the quantification of differential gene expression.

## MATERIALS AND METHODS

### The EmpiReS approach and method

#### EmpiRe applied to RNA-seq data

MS-EmpiRe (21) is based on an **Empi**rical and **Re**plicate-based, approach, further called EmpiRe, which does not assume a given statistical distribution to detect differential proteins between groups of samples in mass spectrometry proteomics data. These groups can be defined by the user and usually represent replicates for different phenotypes or other experimental conditions. Noise in the measurements and biological variability have to be accounted for to detect objects that vary more than expected by chance. EmpiRe does so based on empirical (log2) error distributions (EEDs) of signals between replicate measurements. For each biological object e.g. a gene, transcript or protein, we compute the fold change between all pairs of replicates (see Figure 1 C). It is crucial that only signals for the same object are compared such that eventual biases cancel out as much as possible. After an appropriate normalization (22, 23) that resolves library size differences by estimating per-sample scaling factors (similar to Zien et al. (24), described in more detail in Ammar et al. (21)), the expected value of the distribution of these errors is zero. How strong fold changes of objects deviate from this expected value is determined by their underlying variance. For mass spectrometry as well as sequencing data, it has been observed that this variance depends on the strength of the signal intensity or read count. To express this correlation, EmpiRe computes many error sub-distributions, each of which is based on a subset of objects that share similar overall expression strength. However, we observed that even within an experiment the amount of noise can differ between conditions. We thus modified EmpiRe to use a specific empirical error distribution for each condition consisting of a group of samples.

**Figure 1.**
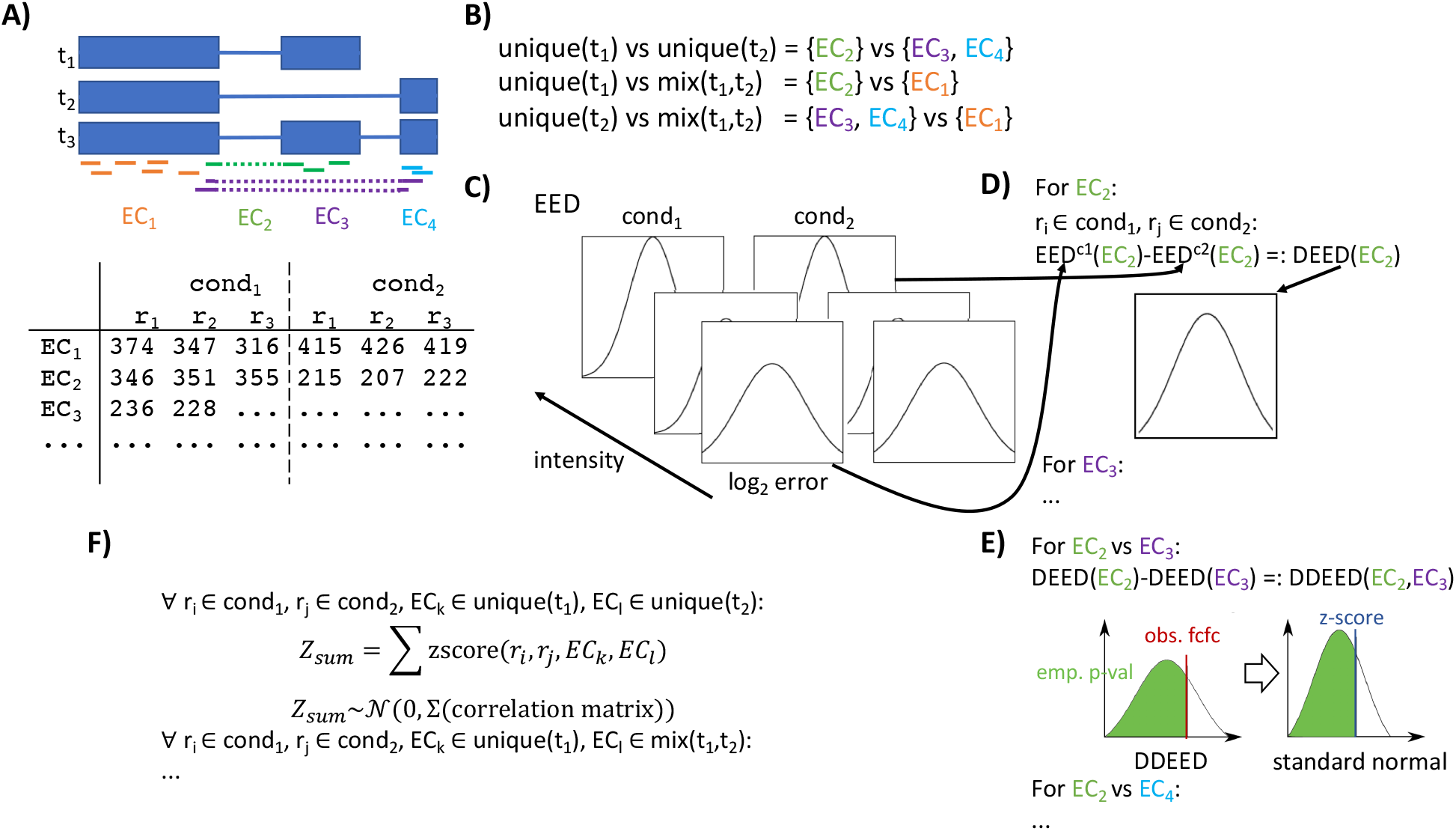
EmpiReS in a differential splicing analysis to quantify transcript expression changes. A) Given the annotation and mapped reads, equivalence classes (ECs) are derived by determining the set of transcripts that are in agreement with a read. For each EC the reads are counted for all replicates for both conditions. B) On the other To test differential alternative splicing (DAS) between two transcripts, we compare the ECs that are either unique for one of the two transcripts or that are a mixture of both transcripts. If any of the three comparisons is significant we classify this as a DAS event. C) For each condition we calculate all within condition fold changes between replicates to derive empirical error distributions (EEDs) for different intensity bins. D) For a given EC and pair of replicates from the two conditions, we select the corresponding EEDs and derive the differential empirical error distribution (DEED) by subtracting the EEDs and assuming independence. E) Similarly, for two ECs we can calculate the error distribution of the change of changes between the ECs, i.e. the doubly differential error distribution (DDEED) by subtracting the corresponding DEEDs. The observed change of changes for the pair of ECs and the pair of replicates can be transformed into a Z-score using the modified Stouffer’s method used in MS-EmpiRe. F) The Z-scores for all combinations of replicates and ECs of the corresponding DAS test are calculated and summed up. The resulting sum follows a 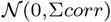 distribution as each individual Z-score is from 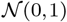

The EEDs are used to assess the significance of an observed fold change for an object between two samples from different conditions *c*_1_ and *c*_2_. For this, we derive the differential empirical error distribution (DEED), which is simply 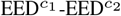, as we can assume that observations from 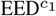 and 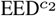 are independent as they come from different measurements (see Supplement section 3 for further details). The significance of an observed fold change can directly be expressed as an empirical *p*-value which estimates the probability to observe a fold change of this strength only due to errors. These empirical *p*-values however do not directly include information about consistency among replicate data or the direction of change. Does a gene change in all comparisons between the conditions and if so, is this change of similar strength and, in particular, in the same direction? To combine *p*-values over multiple comparisons as well as sub-measurements, e.g. peptides, EmpiRe applies an approach similar to Stouffer’s method (25). First, the empirical *p*-values for all replicate comparisons are transformed into signed *Z*-scores (Figure 1 E) that express the same cumulative probability. This is done by using the inverse cumulative distribution function of the standard normal distribution. To keep the direction of the fold change, we assign its sign to the *Z*-score. In order to aggregate these *Z*-scores into one score per object, EmpiRe sums them up to compute an aggregated value, *Z*_*sum*_. Since the *Z*-scores are signed, only objects that show multiple deviations in the same direction become significant. Since each random variable of the individual *Z*-scores follows a normal distribution with mean 0 and variance 1 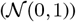, we can also compute the distribution of *Z*_*sum*_. The resulting distribution is normal with mean zero. The variance is equal to the sum over the covariance matrix of the *Z*-scores. The original Stouffer test is applicable to independent random variables which means that this variance is equal to the number of *p*-values combined. We have to account for the fact that two fold changes (and therefore the corresponding *Z*-scores) that share either the first or second signal for the fold change computation are not independent. For all other *Z*-score pairs, the covariance is zero. The derivation of the covariance of two dependent *Z*-scores can be done analytically and the total variance of *Z*_*sum*_ depends only on the number of dependent replicate comparisons and the variances of 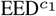 and 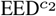 (see Supplement section 6).

EmpiRe makes no data specific assumptions besides that noise of the measurement can be estimated by signal changes between replicate data. The error distributions are derived empirically for the dataset at hand. In Ammar et al. (21) we showed that EmpiRe is applicable to a wide range of mass spectrometry proteomics setups. Here, we demonstrate that the approach incorporating condition-dependent EEDs is also applicable to RNA-seq count data. Furthermore, we show that the approach can be generalized such that it can be applied to doubly differential cases like DAS.

#### Object definition and transcript level counts

In DE analyses, signal measurements (feature values) such as microarray probe set levels, read counts or peptide intensities are often aggregated to obtain an object value, i.e. feature values for genes or proteins. Due to the vastly differing ranges of the raw signals this summarization step and, thus, also the resulting object fold changes are very unreliable. Summarizing raw signal values can be robust when the respective values are quite similar or when very many values are summarized, in general signal summarization should be avoided.

Therefore, Erhard and Zimmer (26) proposed to use local fold changes of measured features (probe sets, read regions, or peptides) between conditions and summarize these local fold changes to obtain respective object fold changes. To analyze differential alternative splicing the summarization problem is aggrevated as the non-overlapping regions between the transcripts can be small and, thus, only few signals are available to estimate the difference between the object (i.e. transcript) feature values. Thus, we propose to directly estimate the fold change between the transcripts from the local signal fold changes and avoid estimating transcript levels altogether.

Available tools can be divided in two major groups depending on how they obtain and use transcript level counts or similar proxies (27): they either directly estimate abundances on the transcript level (Cufflinks (28)) or rely on counts of smaller objects like exon bins (DEXSeq (19)), individual splice junctions (rMATS (20)) or equivalence classes (BANDITS (29)). The first group of models tries to assign each read to its transcript of origin. If a certain splice junction or exon is part of only one transcript and is covered by a read, this assignment is trivial. Reads that could have been generated by different transcripts are assigned by using statistical methods like the Expectation-Maximization algorithm. In contrast, count based models either completely ignore ambiguous reads or assign them to multiple objects.

Probabilistic read assignment as well as multi-assignment may result in errors which are then propagated to the differential analysis. hand, information is lost by neglecting ambiguous reads. Therefore, we decided to use equivalence classes (EC), similar to BANDITS, for EmpiReS: An equivalence class is a set of transcripts with which a set of reads is compatible and they are, thus, constructed by deriving for each read the set of transcripts that are compatible with it (see Figure 1 (A)). However, for genes with many annotated transcripts this can lead to a large number of ECs that may each only contain a small number of reads, even though only few of the annotated transcripts are really expressed. In this case we propose to reduce the number of ECs by a heuristic clustering of the transcripts (see Supplement section 4). Pairs of transcripts with only few reads that discriminate between the two transcripts are merged until a given number of transcript clusters is reached. Then, the ECs are determined using the clustered transcript definitions, which will yield an equal or lower number of ECs for which counts are derived.

As EmpiReS works at the level of ECs we can use a mapper that uses the transcript annotation and the concept of ECs to speed up the mapping and remove ambiguities. A simple modification of contextmap (30) which works at the level of ECs is used as the default mapper for EmpiReS (see Supplement section 5 on method details).

#### Extension to doubly differential setups

The problem of detecting DAS can be generalized: the goal is to detect whether changes of changes of objects differ significantly. In case of DAS, the objects are transcripts or smaller sub-units like exon bins or equivalence classes which do not overlap between transcripts. Instead of looking at the expression change of one object (e.g. gene) we want to know whether the expression of two objects (transcripts of the same gene) changes differently. To allow EmpiReS to be applied to cases like this, we propose the following extensions:

For any possible (clustered) transcript pair (*t*_1_*,t*_2_) of some gene, EmpiReS compares three sets of ECs (see Figure 1 (B)): ECs that are unique for *t*_1_ and *t*_2_ (unique(*t*_1_) and unique(*t*_2_), respectively) and ECs that are a mixture of *t*_1_ and *t*_2_ (mix(*t*_1_, *t*_2_)). For each pair of these sets we determine whether the corresponding changes of changes are significant using the extended model of EmpiReS, and call the two transcripts changing if any of the three comparisons is significant.

Analogously to the approach for differential expression in RNA-seq data we first define the empirical error distribution (EED) for each condition. For a pair of replicates from the two conditions respectively, we select for each EC the EEDs according to the intensity levels of the EC in the two replicates and derive the differential empirical error distribution (DEED) (see Figure 1 (C) and (D)). Similarly, we can derive the doubly differential empirical error distribution (DDEED) by combining the DEEDs of two ECs that we want to compare (see Figure 1(E)). The DDEED is the empirical error distribution of a change of changes between two ECs (i.e. features of different isoforms) between two replicates of different conditions. It can thus be used to determine an empirical p-value for the observed fold change of fold changes for this combination of ECs and replicates. This p-value indicates how likely it is to observe a value that is at least as extreme only due to biases. The p-value is again transformed into a signed Z-score.

The Z-scores of all combinations of replicates and ECs are computed this way and summed up. As each individual Z-score follows a normal distribution 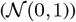, the sum also follows a normal distribution with mean 0 and the sum of (the full covariance matrix of the individual Z-scores as variance. As some of the individual Z-scores share a replicate or EC, they are not independent and the corresponding entry of the covariance matrix will not be zero. See the Supplement section 7 for a detailed description of how the covariance matrix of the individual Z-scores is calculated.

We repeat this step for all pairs of EC sets shown in Figure 1 (B). Finally, p-values are computed as the probability that the observed *Z*_*sum*_ value is expected under the null hypotheses of no differences among replicate measurements. To obtain a single gene level score, we select the highest p-value of the different EC set combinations since a difference between any of these is an indicator for differential splicing.

### Simulation and Benchmark

To simulate reads for the evaluation of differential expression and alternative splicing often a statistical distribution is parameterized and random counts are sampled from this distribution and random (uniform) read positions are derived for these counts. As the different established methods use different types of statistical distributions in their model this way to simulate a benchmark dataset intrinsically favors the methods that use the same distribution. Therefore, we use a simulation approach that directly uses the distribution of counts and read positions from real experiments. Especially for the simulation of DAS events, there are three difficulties that have to be addressed: (i) the splicing event has to be detectable, i.e. there have to be enough reads from the affected region (ii) replicate measurements have to be generated in a way that accurately reflects the variation of replicates in real measurements and (iii) the position-wise distribution of the reads over a transcript has to reflect the biases observed in real data.

To address the detectability of simulated splicing events we select pairs of transcripts inducing DAS-events with long enough differences (i.e. many base pairs differ between transcripts). As our selected source of simulation (human Ensembl, GRCh37.75) has a rather complex transcript annotation we choose to simulate clear simple cases: two transcripts implying at least one exon skipping event. For this we derive all exon-skipping events from the input annotation and select the transcript pair from the corresponding genes where the implied number of skipped bases is maximal.

Replicate measurements are usually simulated by parameterizing a distribution (e.g. a negative binominal distribution) and drawing counts from it. Fold changes are usually produced by applying a multiplicative factor to the sampled counts or sampling with a different parameter. Such simulations, thus, need to make assumptions about the underlying distribution and change the library sizes by scaling simulated counts.

Here, we propose an approach that directly uses real measurements and does not affect library sizes. To simulate a benchmark dataset for differential expression we use gene-level counts of a real experiment with a large number of replicates and divide the replicates of one condition into two groups (the simulated conditions). This way we have for both simulated conditions replicates that differ only by technical and biological biases. To introduce changing genes between the simulated conditions we first select genes from the whole signal strength spectrum by selecting genes from regularly spaced ranks of the sorted list. For each selected gene we sample a target fold change 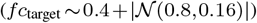 and then select the target gene whose mean signal is closest to the signal that the target fold change implies. The parameters for the target fold change were selected such that the minimal observable change will be 0.4 and 97% of the changes will be less than 4-fold. Finally, we swap the counts of the selected and target genes in one of the simulated conditions to generate target fold changes for the selected genes and the reciprocal fold changes for the target genes without changing the library size.

A popular way to simulate DAS is to distribute gene level counts among a subset of transcripts (31, 32). DAS signals can then be simulated by changing the proportions of this assignment between conditions. While we also re-distribute gene level counts, we use a similar swapping procedure as for the simulation of differential genes to simulate DAS events. In this case we need two genes for both transcripts (major and minor isoform) each that are swapped to simulate a fold change both between the conditions for both isoforms and a fold change between the two isoforms. See the Supplement section 8 for a detailed description of how the swapping partners are selected, in short we first sample a base fold change for the major isoform, that can also be zero, and select the swapping partner of the major isoform. Then another fold change is sampled for the minor isoform and the swapping partner of the minor isoform is selected by searching for a transcript with a mean signal closest to the signal that corresponds to the sum of the two fold changes.

So far we have simulated counts of transcripts containing both differential genes and differential alternative splicing and replicates, but ultimately, we need to simulate reads corresponding to these counts. As we do not want to make any assumptions about the position-wise biases of reads, we use real yeast RNA-seq data to derive start/end fragment positions - ideally there should be virtually no splicing in annotated transcripts in yeast. These derived position-wise biases are mapped and scaled for human transcripts selected for simulation. This means for a human transcript of transcribed length *l* a yeast template transcript will be searched with similar length *l*_*t*_. If the template transcript is longer the subsequence of length *l* with the most observed start/end reads is used. If the template transcript is shorter (*l*_*t*_ < *l*) then *l* − *l*_*t*_ randomly selected positions are assigned a probability of zero. Using these derived position-wise probabilities of the fragment start positions, the start positions are sampled. To determine the corresponding fragment end positions, the insertion length is sampled from a 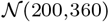 distribution, and the end position is sampled similarly to the start position in a window of size 5 around start+insertion size.

The scripts for the simulation that can be used to simulate data based on any dataset with a sufficient number of replicate measurements, can be downloaded from https://www.bio.ifi.lmu.de/software/empires/index.html. For the evaluation of the DE and DAS methods we used three different human RNA-seq datasets (ECTO, STEM and EBV) with different numbers of replicates and sequencing depth. Suppl. Table 1 contains a short overview of the used datasets. We applied the DE methods DESeq, edgeR, limma, MS-EmpiRe as well as the DAS methods BANDITS, DEXSeq, DRIMSeq, rMATS and our own method to the simulated datasets. For details on how the data was processed and how the methods were called see the Supplement section 2.

## RESULTS

### Evaluation of differential expression analysis

We evaluate the performance of EmpiRe on simulated data for differential gene expression. Due to its successful application to peptide intensities of mass spectrometry data in Ammar et al. (21), we want to demonstrate its applicability to count data from RNA-seq. Multiple simulations were created using different datasets as input for the simulation, differing in the number of replicates and sequencing depth. Simulated genes with too low read counts across samples (as defined by the filterByExpr function of edgeR) are filtered, which results in ≈ 9.500 to 12.500 genes. The MS-EmpiRe approach can directly be applied to read count data, so we evaluate the performance of MS-EmpiRe and its modified approach EmpiRe which includes condition-specific EEDs. Additionally, we apply three established differential expression detection tools: DESeq2 (33, 34), edgeR (35) and limma (36). While the former two are specifically designed for read count data, limma is a more general model, originally developed for RNA microarrays.

Table 1 shows the results of the performance evaluation for differential expression methods. Since the differences in performance between the tools are marginal, we do not report and discuss them in detail. Both auroc and auprc are above 99% for all methods in all simulations indicating a close to perfect separation between differential and non-differential genes. If we look at the precision and recall, differences between the tools become larger. Both measures report the performance at a certain score cutoff. A common choice is to use the multiple testing corrected *p*-value for each gene, further denoted *P_adj_* or FDR. Precision reports how many of the detected genes are actually differential while the recall measures how many of all differential genes were detected. We typically apply a FDR cutoff of 5%, which is quite common since it means that the expected precision at this cutoff is 95%. High precision is important since type I errors, i.e. falsely detecting a non-differential gene, often have a greater impact on downstream analyses or follow-up experiments than missing a few of the differential genes. Not all of the tools achieve a precision of at least the expected value of 95% at about the same high recall value. EdgeR and limma only achieve a precision of about 84% for the simulation based on the STEM dataset, and for the EBV simulation all methods yield precision values below 95%. While EmpiRe is only slightly below 95%, MS-EmpiRe and DESeq have precision values of around 90% and edgeR and limma reach even only about 80%. Apparently, even though the ordering of the genes is nearly perfect, the distribution assumption free approach of EmpiRe is better able to capture the variance and control FDR than the established tools.

**Table 1.** Evaluation results of the differential expression methods. While all methods perform very well in terms of auroc and auprc, EmpiRe achieves the best precision and, thus, nearly achieves the FDR cutoff of 5% also for the more challenging EBV dataset.

|  | method | auprc | auroc | f1 | prec. / recall | TP / FP |
| --- | --- | --- | --- | --- | --- | --- |
| STEM | DESeq | 99.71 | 99.91 | 98.59 | 97.46 / 99.74 | 772 / 20 |
|  | edgeR | 99.67 | 99.88 | 91.35 | 84.27 / 99.74 | 772 / 144 |
|  | limma | 99.69 | 99.91 | 91.41 | 84.37 / 99.74 | 772 / 143 |
|  | MS-Empire | 99.64 | 99.97 | 98.59 | 97.46 / 99.74 | 772 / 20 |
|  | Empire | 99.63 | 99.83 | 99.03 | 98.59 / 99.48 | 768 / 15 |
| EBV | DESeq | 99.86 | 99.99 | 93.65 | 88.05 / 100.00 | 774 / 105 |
|  | edgeR | 99.82 | 99.99 | 89.48 | 80.96 / 100.00 | 774 / 182 |
|  | limma | 99.86 | 99.99 | 89.17 | 80.46 / 100.00 | 774 / 188 |
|  | MS-Empire | 99.74 | 99.97 | 94.97 | 90.42 / 100.00 | 774 / 82 |
|  | Empire | 99.60 | 99.98 | 97.10 | 94.83 / 99.48 | 770 / 42 |

EmpiRe, although not designed specifically for counts and sequencing data, is able to control the FDR at the desired level while detecting nearly all differential genes.

### Evaluation of differential alternative splicing on simulated data

As EmpiReS performed well on differential gene expression we applied it to our simulated differential alternative splicing data and compared its performance to established tools for this task: BANDITS (29), DEXSeq (37), DRIMSeq (17) and rMATS (20).

Table 2 shows the evaluation results for simulations based on two different datasets. The performance of the DAS methods is in general worse than for DE and there are more differences between the methods. As for DE we applied a FDR cutoff of 5% to the predictions and calculated precision and recall. BANDITS and rMATS are not very sensitive and thus achieve only low recall values. All methods are not able to properly control for the FDR, with DRIMSeq and EmpiReS yielding the best precision for the simulation based on the STEM dataset at 92% and 89%, respectively, and EmpiReS (88%) for the simulation based on the EBV dataset. EmpiReS performs best in terms of f1 measure for both datasets, indicating that it provides the best trade-off between precision and recall. Next, we evaluated the ordering of the prediction by calculating auroc and auprc. Here, auprc is better suited for inbalanced benchmark sets such as ours, where the number of true and false examples differs greatly. Again, DRIMSeq and EmpiReS performed best on the STEM simulation (auprc of 0.87 and 0.84, respectively), while EmpiReS is the only method to achieve a auprc above 0.7 for the EBV simulation.

**Table 2.** Evaluation results for the differential alternative splicing methods using 100bp long reads and biased read positions. Over all datasets, EmpiReS achieves the best results both in terms of f1 measure as well as for auprc.

|  | method | auprc | auroc | f1 | prec. / recall | TP / FP |
| --- | --- | --- | --- | --- | --- | --- |
| STEM | BANDITS | 0.42 | 0.50 | 0.32 | 66.67 / 0.16 | 2 / 1 |
|  | DEXSeq | 0.74 | 0.95 | 70.93 | 81.72 / 62.66 | 787 / 176 |
|  | DRIMSeq | 0.87 | 0.93 | 62.68 | 91.99 / 47.53 | 597 / 52 |
|  | rMATS | 0.61 | 0.79 | 47.18 | 60.45 / 38.69 | 486 / 318 |
|  | EmpireS | 0.84 | 0.92 | 78.02 | 89.83 / 68.95 | 866 / 98 |
| EBV | BANDITS | 0.45 | 0.66 | 28.40 | 72.08 / 17.68 | 222 / 86 |
|  | DEXSeq | 0.52 | 0.94 | 59.70 | 46.81 / 82.40 | 1035 / 1176 |
|  | DRIMSeq | 0.59 | 0.86 | 57.66 | 61.16 / 54.54 | 685 / 435 |
|  | rMATS | 0.53 | 0.69 | 40.11 | 65.52 / 28.90 | 363 / 191 |
|  | EmpireS | 0.71 | 0.90 | 75.26 | 87.97 / 65.76 | 826 / 113 |

Furthermore, we compared the performance of the methods on different simulation runs with different datasets (STEM, ECTO or EBV), read lengths (60 or 100) and position-wise bias (bias taken from yeast data or unbiased that is uniformly distributed).

Figure 2 shows the performance of the methods on biased and unbiased simulations. BANDITS and DEXSeq yield lower recall and precision values for the unbiased simulations, while DRIMSeq and rMATS perform better on the unbiased simulations. DRIMSeq has a higher precision on unbiased data but slightly lower recall, while rMATS achieves a higher recall on unbiased data with slightly lower precision. Only EmpiReS performs similarly on both the simulations with and without position-wise bias with nearly the same precision and recall values. Moreover, the precision and recall values show the least variation between the different simulation setups for EmpiReS. In our interpretation this indicates that approaches like DRIMSeq strongly depend on the assumption of uniform read distribution on the transcript sequence, while EmpiReS does not.

**Figure 2.**
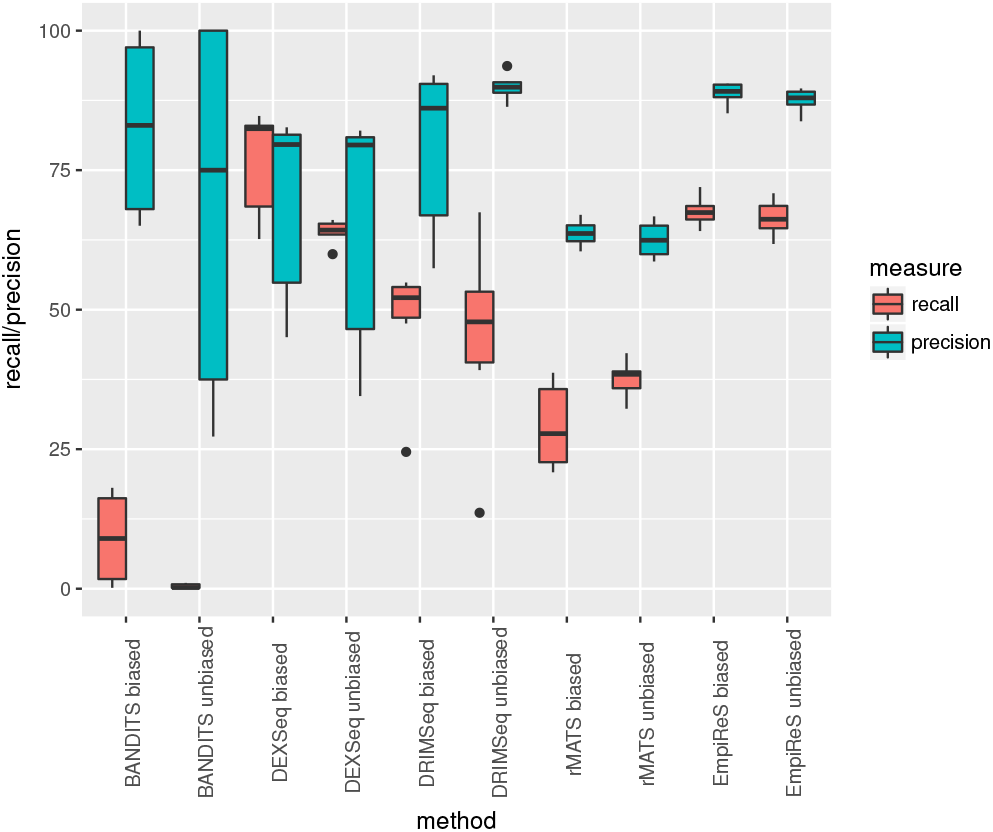
Influence of biased read positions on the performance results for different simulation runs (read lengths 60 or 100, dataset EBV, ECTO or STEM and read positions biased or uniform). Only EmpiReS yields similar precision and recall values for all simulations independent on whether the read positions were biased or not.

### Influence of read mapping on performance of splice event detection

Figure 3 compares the performance of DEXSeq and EmpiReS for different mappings: the mapper that is normally used for this method (HISAT2 for DEXSeq and EC-contextmap for EmpiReS), contextmap and the ideal mapping, where reads are mapped exactly like they were simulated. For DEXSeq the recall is comparable for all mappers at about 60%-80%, while the precision (45%-80%) is slightly better for HISAT2 than for contextmap and best for the ideal mapping. Overall the influence of the mapping on DEXSeq’s performance is modest. For EmpiReS, however, both the recall (60%-70%) and precision (80%-95%) are improved from contextmap to EC-contextmap to the ideal mapping. Especially the precision is improved for all simulation setups and nearly always lies above 90% when the ideal mapping is used showing that errors in the DAS detection originate in the ambiguity of the read sequencing leading to mapping errors and thus to wrong signals.

**Figure 3.**
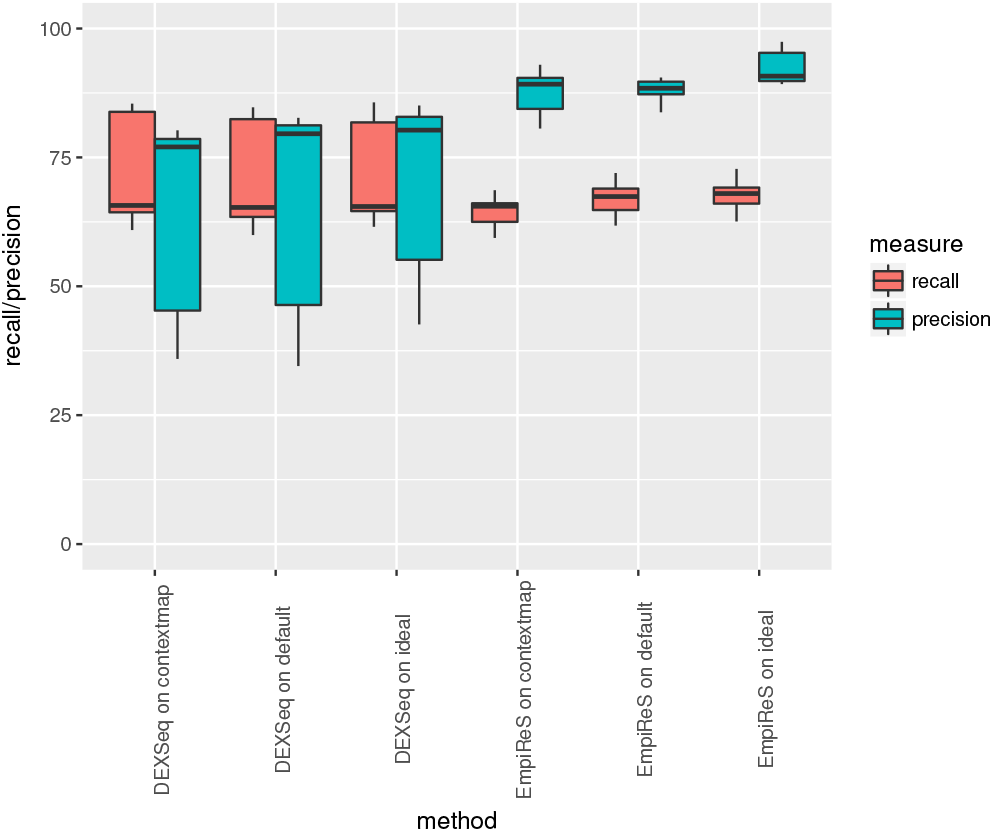
Performance of DEXSeq and EmpiReS on different mappings for different simulation runs. Both methods were applied to mappings of contextmap, their default mapper (HISAT2 for DEXSeq and EC-contextmap for EmpiReS) and the ideal mappings, that is reads are mapped exactly where they were simulated.

### Evaluation of validated DAS events

We used two published datasets for which several differential alternative splicing events have been validated experimentally. Bebee et al. (38) (GEO accesssion GSE64357) compares wild-type and Epithelial splicing regulatory proteins (Esrp) double knocked-out mouse cells. For each condition only 2 replicates were measured and overall 28 differentially spliced events were validated. Shen et al. (20) (SRA accesssion SRP014759) measures two human prostate cancer cell lines with 3 replicates per cell line and validated 32 splicing events.

Table 3 shows how many events were called by the different DAS methods and how many of the validated events were among the called events. For the Shen dataset nearly all methods found all of the validated splicing events, only DEXSeq missed one of the 32 events. However, the methods differ vastly in the number of called events, ranging from 4.100 for rMATS to 8.700 predicted events for DEXSeq. EmpiReS with about 5.200 predicted events is comparatively conservative, but still manages to detect all validated splicing events. For the Bebee dataset there were more differences. The number of predicted splicing events ranged from about 90 for BANDITS to over 1.000 for DEXSeq. Also the number of validated splicing events that were predicted from the DAS methods differed more than for the Shen dataset. No method was able to predict all of the 28 validated splicing events. This is most likely explained by the low statistical power when only two replicates are measured. The two methods that predict the highest number of events (DEXSeq and rMATS) also find the highest number of validated events (23 and 26, respectively). EmpiReS finds 17 of the 28 validated splicing events, while only predicting 520 events in total. Thus, while only predicting half as many events it still manages to find about two thirds of the validated events, indicating a higher precision, just as observed in the simulations.

**Table 3.** Overview of the results of the DAS methods for the validated splicing events for the Bebee and Shen datasets. For the Shen dataset all methods found nearly all of the 32 validated splicing events, while there are greater differences for the Bebee dataset that consists of only two replicates and for which 28 splicing events were validated. DRIMSeq and EmpiReS predict only half as many events as rMATS and DEXSeq, but also misses more of the validated splicing events.

| method | Beebe |  | Shen |  |
| --- | --- | --- | --- | --- |
|  | called | vali. found | called | vali. found |
| BANDITS | 94 | 11 | 4149 | 32 |
| DEXSeq | 1124 | 23 | 8723 | 31 |
| DRIMSeq | 649 | 18 | 5978 | 32 |
| rMATS | 974 | 26 | 4121 | 32 |
| EmpiReS | 520 | 17 | 5186 | 32 |

## DISCUSSION

The increasing number of high throughput measurement techniques makes differential analysis methods and pipelines for omics data ever more important. A common analysis step is the differential quantification of certain features of measured objects across conditions or compared to controls. Typical are differential gene expression and differential alternative splicing. Consequently, there are many methods and associated tools available for these purposes. But as the results depend quite intricately on the applied methods, the user needs to have detailed knowledge about the models employed in the respective methods and, even then, it is unclear how to deal with inconsistent and contradicting results. In order to assess the available tools it is preferable to detect as many true and relevant events (true positives) as possible while keeping the number of false discoveries as small as possible. Typically, the user specifies a FDR threshold, e.g. 5%, such that it is expected to obtain only five false predictions among every hundred positive predictions of the method.

With MS-EmpiRe, we have previously introduced an approach for the quantification of differential proteins in mass spectrometry data based on measured peptide intensities. The EmpiRe method introduced here shows how to exploit the EmpiRe approach for differential gene expression for read count data derived from RNA-seq measurements. Moreover, we generalize the approach for doubly differential setups.

To evaluate both differential expression and differential alternative splicing, we use a simulation approach that makes minimal assumptions. Instead of sampling reads from some predefined and assumed distribution, we use measurements from real experiments and introduce differential genes or alternative splicing events by swapping the labels of genes/isoforms in a subset of replicate samples. This way the distribution of the counts of the real experiment is retained. Moreover, the position of the reads can also be distributed according to a real experiment in yeast where no biases caused by splicing events should be present. The resulting simulated datasets are thus more realistic as they retain all biases present in a given real experiment. This simulation approach can be applied to any real experiment to be used in future benchmarking studies.

EmpiRe yields results comparable to established tools for differential gene expression such as DESeq, edgeR and limma, but significantly improves the precision. It thus predicts less false positives compared to the established methods.

Additionally, we show how to extend EmpiRe to doubly differential analyses (EmpiReS) and report results on the quantification of differential alternative splicing (i.e. a typical doubly differential setup). Compared to the state of the art tools DEXSeq, BANDITS, DRIMSeq, and rMATS, EmpiReS yields overall better trade-offs between precision and recall for different simulation setups using different sequencing depths and biases derived from real datasets. Moreover, its performance was most robust with respect to different simulation setups and did not depend on whether or not a position-wise bias was present in the data.

EmpiRe(S) offers a general unified approach to differential analyses. We have shown that its lack of assumptions makes it applicable to various data types (mass spectrometry derived intensities, sequencing based read counts) and non-trivial research questions such as differential alternative splicing. Since EmpiRe(S) depends on replicated measurements to allow for empirical estimation of measurement errors and biological variation, it requires at least two replicates per condition. We think that EmpiReS will also be useful for more complicated setups such as time series measurements over multiple conditions, population studies with many individuals, and in particular single cell measurements, even more so as many more (biological) replicates will be available to estimate the required error fold change distributions very accurately.

## FUNDING

This work has partially been supported by the DFG via the CRC 1123 Atherosclerosis.

## Notes

https://www.bio.ifi.lmu.de/software/empires/index.html

### Competing Interest Statement

The authors have declared no competing interest.

