## Supplement for "EmpiReS: Differential Analysis of Gene Expression and Alternative Splicing"

August 4, 2020

#### 1 Data

As basis for our simulation we selected two datasets from the Human Reference Epigenome Mapping Project for HUES64 cells (STEM) and CD56+ ectoderm cells (ECTO). These datasets have many replicates (see Table 1) and are fairly deep sequenced (see Figure 1). Additionally, we also created a dataset of much deeper sequenced replicates by pooling together replicates of different conditions of human B cell line samples used for analysis of Epstein-Barr virus infection (EBV). This dataset consists of 10 different conditions with 3 biological replicates with 2 technical replicates each (i.e. 6 replicates per condition). We generated 6 pooled replicates by combining replicates of different conditions to generate a very deeply sequenced dataset. The pooled replicates are created in pairs so that the two technical replicates for each biological replicate are always added to the same pair of pooled replicates. In the subsequent simulation these pairs of pooled replicates are assigned to the two simulated conditions, thus ensuring minimal variability between the conditions before the swapping procedure to add differential genes/isoforms starts.

| name | # replicates | SRA ids |
|---|---|---|
| ECTO | 13 | SRR1067050, SRR1067052, SRR1067053, SRR1067054, SRR1067056, SRR1067057, SRR1097443, SRR1097445, SRR1097446, SRR1097447, SRR1097448, SRR1097449, SRR1097450 |
| STEM | 6 | SRR1067487, SRR1067488, SRR1067492, SRR1107846, SRR1107848, SRR1107849 |
| EBV | 60 pooled to 6 | - |

Table 1: Overview of the human datasets used to simulate counts for the evaluation.

To simulate the position-wise bias we used a dataset measuring the yeast heat shock response with overall 60 samples. All these samples are pooled together to achieve a very high coverage. As there is no alternative splicing in yeast we can use this dataset to simulate the position-wise bias for human transcripts as described in the paper.

#### 2 Related Methods

For the evaluation of the differential expression methods we applied the methods to the simulated counts. The EnrichmentBrowser (Geistlinger *et al.*, 2016) was used to call DESeq, limma and edgeR with default parameters.

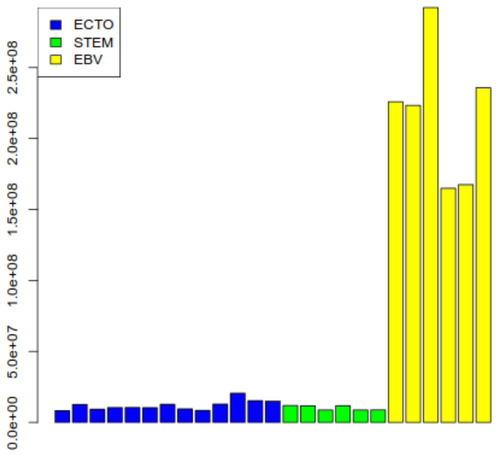


Figure 1: Sequencing depths for the human data sets used for simulation. The mean sequencing depth is 11.98 Mio reads for ECTO (blue), 10.27 Mio reads for STEM (green) and 218.28 Mio reads for EBV (yellow).

Each alternative splicing method was called with reads mapped with its preferred mappers: kallisto for DRIMSeq and BANDITS, HISAT2 for DEXSeq and rMATS. Kallisto (version 0.45.0) is called with 100 bootstrap samples and bias correction. For HISAT2 (version 2.1.0) the downstream transcript assembler option was used. DEXSeq and rMATS (version 4.0.2) were called with standard parameters. For DRIMSeq the dmFilter method used to filter genes and features with low expression was called with the parameters `min_samps_gene_expr=#samples, min_samps_feature_expr=3, min_gene_expr=10, min_feature_expr=10` and for BANDITS the filter_transcripts method to filter transcripts with low expression was called with the parameters `min_transcript_proportion=0.01, min_transcript_counts=10,min_gene_counts=20`.

### 3 Combining empirical distributions

To derive the differential empirical error distribution (DEED) we combine 2 empirical error distributions (EEDs). This corresponds to defining a new random variable $D := R_1 - R_2$ from the random variables $R_1$ and $R_2$ that correspond to the EEDs that should be combined. The empirical distributions are saved as relative frequencies for (small) bins of the range of all observed values. For each combination of bins of the two EEDs the value (i.e. bin) of the combined distribution can be calculated by simple subtraction. The relative frequency of this new value in the new empirical distribution is defined by the probability of observing the values from the EEDs at the same time, which can be calculated by the product of the individual relative frequencies, given independence of the two EEDs. As the EEDs are derived from independent measurements, the assumption of independence is valid. As multiple combinations of values from the two EEDs can yield the same fold change in the DEED the relative frequency of a fold change is the sum of the relative frequencies of all combinations yielding this fold change value:

$$P(D(fc)) = P(\bigvee_{\forall s_1,s_2: fc=s_1-s_2} R_1(s_1) \wedge R_2(s_2)) = \sum_{\forall s_1,s_2: fc=s_1-s_2} P(R_1(s_1)) * P(R_2(s_2)) \quad (1)$$

#### 4 Clustering to reduce Equivalence Classes

Even a small number of expressed transcripts can lead to a high number of observed ECs due to the complex transcriptome annotation. Most of the ECs however have only very few reads assigned and are not really useful for any test, and moreover, using all combinations would lead to a high number of tests which would hamper the runtime of the analysis.

Based on the assumption that in most real data there is a rather limited set of expressed isoforms per gene we designed a simple heuristic procedure that derives the most likely set of expressed transcripts from the total set of observed ECs with their corresponding counts. The output is a restricted set of transcripts (or rather clustered transcripts), which in turn are used in a second iteration to derive a new reduced set of ECs which is used for the DAS tests.

The procedure is a simple greedy heuristic: we start with the singleton clusters: every transcript belongs to a cluster only containing the transcript. Then in each iteration we select a pair of clusters to be merged based on the implied ECs and observed counts per EC. The score used in the selection of the next clustering step is a simple heuristic one defined for a pair of transcripts derived from the observed total counts (summed up from all samples used for the analysis):

$$\text{score}(t_1, t_2) = \log_{10}((t_1 \backslash t_2 \text{ counts}) + 2) * \log_{10}((t_2 \backslash t_1 \text{ counts}) + 2) \tag{2}$$

where x\y counts refers to the number of read(-pairs) mapped to x, but not to y. For every transcript, we collect all pairwise scores with all other transcripts with overlapping reads, and select the minimal score as transcript-specific score. In every iteration the pair of transcripts $t_1$, $t_2$ with minimal transcript-specific score is selected for clustering and the two clusters containing $t_1$ and $t_2$ are merged.

After the merging of the clusters new ECs are derived - simply by replacing all occurrences of any transcript id from the clusters by the new cluster id. This will always generate less or as many ECs as in the step before (see Figure 2 for a small example). The procedure terminates if the specified number of clusters is reached.

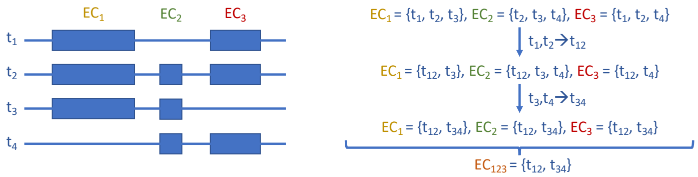


Figure 2: Simple example of the clustering done to reduce the number of ECs. On the left four transcripts and their implied ECs are shown. On the right, two clustering steps and the resulting ECs are shown. In the first step, $t_1$ and $t_2$ yield the minimal score and are merged to $t_{12}$. This does not change the overall number of ECs. In the second step, $t_3$ and $t_4$ are merged to $t_{34}$, after which all three ECs contain the same transcripts so that only one EC ($EC_{123}$) exists after this clustering step.

#### 5 EC-Contextmap

As we derive ECs based on the transcript annotation and only consider ECs for EmpiReS, we developed a fast and simple modification of contextmap that creates a EC-based transcript mapping. For this, we build a suffix array along with LCP-tables based on the transcript sequences extracted from the annotation (GTF-file) and the corresponding genome build. For the mapping we look for longest matching subsequences with at least 10 bases in the reads - i.e. first we look for the longest matching sequence starting from the first base of the read until either all bases are matched or a mismatch occurs at position i, than we start over the matching from the position i+1. For all matched read(-pair)s we keep a table of matched transcript start positions with the number of total matched bases. We use a two-step heuristic filter on all the generated matches: 1) for each read(-pair) we determine the highest number of matches and discard all matching transcript start positions with fewer matches than 80% of this maximal number of matches 2) for every candidate transcript and read(-pair) only the position with the minimal number of mismatches is kept and read(-pair) matchings exceeding 10 mismatches are discarded.

Next, we need to resolve to which gene the read(-pairs) map. All transcript candidate matches are used to create a gene-to-minimal-mismatch mapping. If this results in a single best gene (i.e. the minimal mismatch is achieved with a single gene) then the mapping is resolved and we also update the number of resolved reads for this gene (using all samples of the experiment at the same time). If the minimum mismatch was gene-ambiguous we keep the read(-pair) as ambiguous and will resolve it after we first processed all reads.

After processing all read(-pairs) in the first round we reconsider all gene-ambiguous read(-pairs) and resolve them by using the gene-to-minimal-mismatch mapping using an ad-hoc heuristic. We consider the two genes that the read maps to with the highest number of resolved reads: if the gene with the highest count of resolved reads has at least four times as many resolved mapped reads as the second one, the first one is selected, otherwise the whole mapping is discarded. We do not need to resolve read(-pair)s to individual transcripts, but can directly report the counts for the ECs which are used for EmpiReS.

### 6 Variance of $Z_{sum}$ for DE

If all $Z_i$ in $Z_{sum} = \Sigma_1^n Z_i$ were independent, $Z_{sum}$ would follow $\mathcal{N}(0, n)$ as each $Z_i$ follows a $\mathcal{N}(0, 1)$. However, the $Z_i$ values for fold changes that share a replicate are dependent. The variance of a sum of random variables with dependencies can be calculated by the sum of the covariance matrix. As the correlation is the normalized version of the covariance matrix, we can calculate the variance of $Z_{sum}$ by the sum of the correlation matrix of the corresponding DEED distributions.

To calculate the covariance of two DEED distributions (i.e. entries that are not on the diagonal of the covariance matrix) we have:

$$F_1 := R_i^{c_1} - R_j^{c_2} \qquad R_i^{c_1} \sim EED^{c_1}, R_j^{c_2} \sim EED^{c_2}$$

$$F_2 := R_k^{c_1} - R_l^{c_2} \qquad R_k^{c_1} \sim EED^{c_1}, R_l^{c_2} \sim EED^{c_2}$$

$F_1$ and $F_2$ are the random variables of the errors of fold changes derived from replicates $i$ and $j$ or $k$ and $l$, respectively. They are derived from the random variables of the errors of the corresponding replicate measurements $R_i^{c_1}, R_j^{c_2}, R_k^{c_1}$ and $R_l^{c_2}$

$$\begin{aligned}
Cov(F_1, F_2) &= E[(F_1 - E(F_1)) * (F_2 - E(F_2))] = && E(F_1) = E(F_2) = 0 \\
&= E[(R_i^{c_1} - R_j^{c_2}) * (R_k^{c_1} - R_l^{c_2})] = \\
&= E[R_i^{c_1} R_k^{c_1} - R_j^{c_2} R_k^{c_1} - R_i^{c_1} R_l^{c_2} + R_j^{c_2} R_l^{c_2}] = \\
&= E[R_i^{c_1} R_k^{c_1}] - E[R_j^{c_2} R_k^{c_1}] - E[R_i^{c_1} R_l^{c_2}] + E[R_j^{c_2} R_l^{c_2}] = && \text{exp. value of sum of rand. var.} \\
&= E[R_i^{c_1} R_k^{c_1}] - E[R_j^{c_2}]E[R_k^{c_1}] - E[R_i^{c_1}]E[R_l^{c_2}] + E[R_j^{c_2} R_l^{c_2}] = && \text{prod. of i.i.d. rand. var.}, E[R] = 0 \\
&= E[R_i^{c_1} R_k^{c_1}] + E[R_j^{c_2} R_l^{c_2}]
\end{aligned}$$

in case $R_i^{c_1} \neq R_k^{c_1}$ and $R_j^{c_2} \neq R_l^{c_2}$:

$$\begin{aligned}
&= E[R_i^{c_1}]E[R_k^{c_1}] + E[R_j^{c_2}]E[R_l^{c_2}] && R_j^{c_2}, R_l^{c_2} \text{ i.i.d and } R_i^{c_1}, R_k^{c_1} \text{ i.i.d} \\
&= 0 && E[R] = 0
\end{aligned}$$

in case $R_i^{c_1} = R_k^{c_1}$ :

$$\begin{aligned}
&= E[R_i^{c_1} R_i^{c_1}] + E[R_j^{c_2} R_l^{c_2}] && R_j^{c_2} \neq R_l^{c_2} \text{and i.i.d., as } F_1 \neq F_2 \\
&= var[EED^{c_1}] + E[R_j^{c_2}]E[R_l^{c_2}] && E[R] = 0 \\
&= var[EED^{c_1}]
\end{aligned}$$

analogously for $R_j^{c_2} = R_l^{c_2}$:

$$= var[EED^{c_2}]$$

In total, for an experiment with $n_1$ and $n_2$ replicates in the two conditions, respectively, we thus have:

$$\sum_{i \neq j} Cov(F_i, F_j) = n_1 n_2 (n_2 - 1) var(EED^{c_1}) + n_1 n_2 (n_1 - 1) var(EED^{c_2})$$

as we have $n_1 n_2$ replicate pairs and for each replicate pair we have $(n_2 - 1)$ dependencies that share the replicate of condition 1 and $(n_1 - 1)$ that share the replicate of condition 2.

The $n_1 * n_2$ entries on the diagonal of the covariance matrix each are $(var(EED^{c_1}) + var(EED^{c_2}))$ so that in total we get:

$$\Sigma Cov(F_i, F_j) = n_1 n_2 (var(EED^{c_1}) + var(EED^{c_2}) + n_1 n_2 (n_2 - 1) var(EED^{c_1}) + n_1 n_2 (n_1 - 1) var(EED^{c_2})$$

$$\begin{aligned}
\Sigma Corr(F_i, F_j) &= \frac{\Sigma Cov(F_i, F_j)}{sd(F_i) sd(F_j)} = \frac{\Sigma Cov(F_i, F_j)}{\sqrt{var(EED^{c_1}) + var(EED^{c_2})}\sqrt{var(EED^{c_1}) + var(EED^{c_2})}} \\
&= \frac{n_1 n_2 (var(EED^{c_1}) + var(EED^{c_2})) + n_1 n_2 (n_2 - 1) var(EED^{c_1}) + n_1 n_2 (n_1 - 1) var(EED^{c_2})}{var(EED^{c_1}) + var(EED^{c_2})}
\end{aligned}$$

Since we use the Stouffer transformed z-score rather than the directly observed fold change values, we have to use the correlation matrix instead of the covariance matrix. Using this formula the variance of $Z_{sum}$ can be calculated using only the number of replicates and variances of the EEDs.

### 7 Variance of $Z_{sum}$ for DAS

The individual $Z_i$ in $Z_{sum}$ for DAS are derived from fold changes of fold changes of two features from two feature groups $f_1$ and $f_2$ (e.g. ECs that are unique for isoform 1 and unique ECs for isoform 2) between two conditions $c_1$ and $c_2$ and the corresponding DDEED distribution.

$$\begin{gathered}
FF_1 := (R_{ia}^{c_1, f_1} - R_{ja}^{c_2, f_1}) - (R_{kb}^{c_1, f_2} - R_{lb}^{c_2, f_2}) \\
R_{ia}^{c_1, f_1} \sim EED_a^{c_1, f_1}, R_{ja}^{c_2, f_1} \sim EED_a^{c_2, f_1}, R_{kb}^{c_1, f_2} \sim EED_b^{c_1, f_2}, R_{lb}^{c_2, f_2} \sim EED_b^{c_2, f_2} \\
FF_2 := (R_{mc}^{c_1, f_1} - R_{nc}^{c_2, f_1}) - (R_{od}^{c_1, f_2} - R_{pd}^{c_2, f_2})) \\
R_{mc}^{c_1, f_1} \sim EED_c^{c_1, f_1}, R_{nc}^{c_2, f_1} \sim EED_c^{c_2, f_1}, R_{od}^{c_1, f_2} \sim EED_d^{c_1, f_2}, R_{pd}^{c_2, f_2} \sim EED_d^{c_2, f_2}
\end{gathered}$$

$FF_1$ and $FF_2$ are the random variables of the errors of fold changes of fold changes. Here, the first index $(i, j, k, l, m, n, o, p)$ corresponds to the replicate and the second $(a, b, c, d)$ to the feature. Analogously to the $Z_{sum}$ for DE we can calculate the variance of the DDEED by the sum over the correlation matrix. First we consider the individual entries in the covariance matrix:

$$Cov(FF_1, FF_2) = E[(FF_1 - E(FF_1)) * (FF_2 - E(FF_2))] \qquad E(FF_1) = E(FF_2) = 0$$
$$= E[((R_{ia}^{c_1,f_1} - R_{ja}^{c_2,f_1}) - (R_{kb}^{c_1,f_2} - R_{lb}^{c_2,f_2})) * ((R_{mc}^{c_1,f_1} - R_{nc}^{c_2,f_1}) - (R_{od}^{c_1,f_2} - R_{pd}^{c_2,f_2}))]$$
$$= E[(R_{ia}^{c_1,f_1} - R_{ja}^{c_2,f_1} - R_{kb}^{c_1,f_2} + R_{lb}^{c_2,f_2}) * (R_{mc}^{c_1,f_1} - R_{nc}^{c_2,f_1} - R_{od}^{c_1,f_2} + R_{pd}^{c_2,f_2})]$$
$$= E[(R_{ia}^{c_1,f_1} R_{mc}^{c_1,f_1} - R_{ia}^{c_1,f_1} R_{nc}^{c_2,f_1} - R_{ia}^{c_1,f_1} R_{od}^{c_1,f_2} + R_{ia}^{c_1,f_1} R_{pd}^{c_2,f_2} - \quad \text{remove terms with i.i.d. variables}$$
$$- R_{ja}^{c_2,f_1} R_{mc}^{c_1,f_1} + R_{ja}^{c_2,f_1} R_{nc}^{c_2,f_1} + R_{ja}^{c_2,f_1} R_{od}^{c_1,f_2} - R_{ja}^{c_2,f_1} R_{pd}^{c_2,f_2} -$$
$$- R_{kb}^{c_1,f_2} R_{mc}^{c_1,f_1} + R_{kb}^{c_1,f_2} R_{nc}^{c_2,f_1} + R_{kb}^{c_1,f_2} R_{od}^{c_1,f_2} - R_{kb}^{c_1,f_2} R_{pd}^{c_2,f_2} +$$
$$+ R_{lb}^{c_2,f_2} R_{mc}^{c_1,f_1} - R_{lb}^{c_2,f_2} R_{nc}^{c_2,f_1} - R_{lb}^{c_2,f_2} R_{od}^{c_1,f_2} + R_{lb}^{c_2,f_2} R_{pd}^{c_2,f_2}]$$
$$= E[R_{ia}^{c_1,f_1} R_{mc}^{c_1,f_1} + R_{ja}^{c_2,f_1} R_{nc}^{c_2,f_1} + R_{kb}^{c_1,f_2} R_{od}^{c_1,f_2} + R_{lb}^{c_2,f_2} R_{pd}^{c_2,f_2}]$$

in general the $R^{c,f}$ are independent measurements, that can only be dependent when the same replicate and feature is used. As replicates and features are disjunct between conditions and feature groups, we can eliminate many terms that contain independent variables in the equation above. There are now several cases, depending on which combination of replicates and features are equal:

1. $a = c \wedge b = d \wedge i = m \wedge j = n \wedge k = o \wedge l = p$ (diagonal elements)

$$E[R_{ia}^{c_1,f_1} R_{ia}^{c_1,f_1} + R_{ja}^{c_2,f_1} R_{ja}^{c_2,f_1} + R_{kb}^{c_1,f_2} R_{kb}^{c_1,f_2} + R_{lb}^{c_2,f_2} R_{lb}^{c_2,f_2}] =$$
$$= var(EED_a^{c_1,f_1}) + var(EED_a^{c_2,f_1}) + var(EED_b^{c_1,f_2}) + var(EED_b^{c_2,f_2})$$

2. $a = c \wedge b \neq d$ (first feature shared)

$$E[R_{ia}^{c_1,f_1} R_{ma}^{c_1,f_1} + R_{ja}^{c_2,f_1} R_{na}^{c_2,f_1} + R_{kb}^{c_1,f_2} R_{od}^{c_1,f_2} + R_{lb}^{c_2,f_2} R_{pd}^{c_2,f_2}] =$$
$$= E[R_{ia}^{c_1,f_1} R_{ma}^{c_1,f_1} + R_{ja}^{c_2,f_1} R_{na}^{c_2,f_1}]$$

depending on which replicates are shared, this solves to $var(EED_a^{c_1,f_1})$, $var(EED_a^{c_2,f_1})$ or the sum of both variances when $i = m$, $j = n$ or both.

3. $a \neq c \wedge b = d$ (second feature shared)

$$E[R_{ia}^{c_1,f_1} R_{ma}^{c_1,f_1} + R_{ja}^{c_2,f_1} R_{na}^{c_2,f_1} + R_{kb}^{c_1,f_2} R_{od}^{c_1,f_2} + R_{lb}^{c_2,f_2} R_{pd}^{c_2,f_2}] =$$
$$= E[R_{kb}^{c_1,f_2} R_{od}^{c_1,f_2} + R_{lb}^{c_2,f_2} R_{pd}^{c_2,f_2}]$$

depending on which replicates are shared, this solves to $var(EED_b^{c_1,f_2})$, $var(EED_b^{c_2,f_2})$ or the sum of both variances when $k = o$, $l = p$ or both.

4. $a = c \wedge b = d$ (both features shared)

$$E[R_{ia}^{c_1,f_1} R_{ma}^{c_1,f_1} + R_{ja}^{c_2,f_1} R_{na}^{c_2,f_1} + R_{kb}^{c_1,f_2} R_{ob}^{c_1,f_2} + R_{lb}^{c_2,f_2} R_{pb}^{c_2,f_2}]$$

depending on which replicates are shared, this solves to the sum of the variances of the EEDs corresponding to the terms for which the replicates are shared.

Since we use the Stouffer transformed z-score rather than the directly observed $fcfc$ values, we have to use the correlation matrix instead of the covariance matrix. So each cell from the covariance matrix is normalized with the product of the standard deviations of the corresponding two distributions:

$$\sqrt{var(EED_a^{c_1,f_1}) + var(EED_a^{c_2,f_1}) + var(EED_b^{c_1,f_2}) + var(EED_b^{c_2,f_2})} *$$
$$\sqrt{var(EED_c^{c_1,f_1}) + var(EED_c^{c_2,f_1}) + var(EED_d^{c_1,f_2}) + var(EED_d^{c_2,f_2})}$$

with the corresponding indices for $a, b, c$ and $d$.

### 8 Simulation of DAS events

The procedure to simulate a DAS event is similar to simulating a differentially expressed gene except that two genes with two transcripts (i.e. major and minor isoforms) are swapped to create a DAS event. We want to cover a wide range of signal strength spectrum but also have to ensure that there is a large enough difference between the simulated major and minor isoforms of a gene. First, we order the gene level measurements of the real experiment by their mean signal strength. To select the signal strength for the minor isoform, we start at the 60% quantile of this distribution and decrease the index by max(2, 0.4 * length/num_transcripts) in each step, where length is the number of measurements in the real experiment and num_transcripts the number of transcripts that should be simulated. For the major isoform we start at the 95% quantile and decrease the index by max(2, 0.2*length/num_transcripts). This way, we distribute the major isoforms evenly between the 95% to 75% quantile and the minor isoform between the 60% and 20% quantiles, but we ensure to always increase the index by at least 2.

In each step, two fold changes are sampled from $0.4 + |\mathcal{N}(0.8, 0.16)|$ - a base fold change that corresponds to the changes between conditions of both isoforms, and a isoform specific fold change by which the two isoforms differ in one of the conditions. The base fold change is only applied in 70% of the cases to create a mixture of DAS events for differential and non-differential genes. The target signal values for both isoforms are calculated by adding the base fold change to the signals of both isoforms, and either adding or subtracting the isoform specific fold change to the minor isoform. The swapping partners of both isoforms are selected to be as close to this target signal value as possible. Signal values that were already selected for swapping are masked from the further procedure, so that they are skipped if they would be selected as index for major/minor isoform or swapping partners. If the best available swapping partner would yield a fold change below 0.4 this swapping event is skipped to ensure that all simulated splicing events are detectable.
